# ER exit sites in *Drosophila* display abundant ER-Golgi vesicles and pearled tubes but no megacarriers

**DOI:** 10.1101/2021.03.09.434528

**Authors:** Ke Yang, Min Liu, Zhi Feng, Marta Rojas, Lingjian Zhou, Hongmei Ke, José Carlos Pastor-Pareja

**Affiliations:** School of Life Sciences, Tsinghua University, Beijing, China; School of Medicine, Tsinghua University, Beijing, China; Tsinghua-Peking Center for Life Sciences, Beijing, China

**Author notes:** These authors contributed equally to this work (Ke Yang, Min Liu). Contact: José Carlos Pastor-Pareja School of Life Sciences, Tsinghua University Medical Science Bldg., D224 Beijing 100084, China http://joselab.life.tsinghua.edu.cn.

**Keywords:** Traffic, secretion, intermediate compartment, ERGIC, Golgi, Tango1

## Abstract

Secretory cargos are collected at ER exit sites (ERES) before transport to the Golgi apparatus. Decades of research have provided many details of the molecular events underlying ER-Golgi exchanges. Essential questions, however, remain about the organization of the ER-Golgi interface in cells and the type of membrane structures mediating traffic from ERES. To investigate these, we used transgenic tagging in *Drosophila* flies, 3D-SIM and FIB-SEM to characterize ERES-Golgi units in collagen-producing fat body, imaginal discs and imaginal discs overexpressing ERES determinant Tango1. We found in front of ERES a pre-cis-Golgi region involved in both anterograde and retrograde transport. This pre-cis-Golgi is continuous with the rest of the Golgi, not a separate intermediate compartment or collection of large carriers, for which we found no evidence. We found, however, many vesicles, as well as pearled tubules connecting ERES and Golgi.

## INTRODUCTION

Secretion is one of the most vital processes in the morphogenesis and physiology of eukaryotic organisms, both unicellular and multicellular. In the early secretory pathway, the exit of protein cargos from the endoplasmic reticulum (ER) takes place at specialized ER regions called ER exit sites (ERES), where proteins destined to be secreted are collected prior to their trafficking to the Golgi apparatus (Bannykh et al., 1996). ER-Golgi cargo transfer is the most regulated step in secretion, requiring localized action at ERES of dozens of membrane budding and fusion regulators. Among these, budding of Golgi-bound membrane carriers at ERES is known to involve the COPII coat complex, a set of proteins highly conserved in eukaryotes (Jensen and Schekman, 2011). In these same ERES regions, in addition, traffic in the reverse Golgi-ER direction through COPI vesicles is thought to concentrate as well (Roy Chowdhury et al., 2020). Despite their highly dynamic underlying nature, live imaging has repeatedly shown that ERES are relatively long-lived, stable entities (daSilva et al., 2004; Hammond and Glick, 2000; Shindiapina and Barlowe, 2010; Westrate et al., 2020). Decades of research have given us a detailed view of the molecular events underlying ER-Golgi exchanges from the genetic and biochemical perspectives (Bard et al., 2006; Barlowe and Miller, 2013; Lee et al., 2004). Essential questions, however, remain about the functional organization of the ER-Golgi interface in living cells and the type of membrane structures effectively mediating ER-Golgi traffic from ERES.

Functional organization of the ER-Golgi interface appears to show striking differences across the evolutionary scale. In fission yeasts, protozoa and plants, Golgi stacks remain disseminated throughout the cytoplasm in close proximity to ERES, forming ERES-Golgi units (Brandizzi and Barlowe, 2013; Glick and Nakano, 2009). Invertebrate animals such as the fruit fly *Drosophila melanogaster* (Kondylis and Rabouille, 2003; Ripoche et al., 1994) and the nematode *Caenorhabditis elegans* (Witte et al., 2011) share this type of organization into discrete ERES-Golgi units. In vertebrates, in contrast, vesicles derived from the dispersed ERES fuse to form an ER-Golgi intermediate compartment (ERGIC), through which cargo transits to a single juxtanuclear Golgi ribbon located next to the centrosome (Appenzeller-Herzog and Hauri, 2006). Importantly, the status of the ERGIC as a stable compartment or a transient collection of carriers is unclear. The prevailing view is that it evolved in the larger-sized vertebrate cells as a specialized organelle that, together with the centralized Golgi, optimizes long-distance communication between the perinuclear ER and the plasma membrane (Brandizzi and Barlowe, 2013). However, this is difficult to reconcile with the high conservation of all machineries that regulate ER-Golgi traffic, from yeast to humans and all animals (Saraste and Marie, 2018). Furthermore, electron microscopy studies of *Drosophila* ERES-Golgi units have reported the existence of pleiomorphic elements in the reduced space between ERES and Golgi (Kondylis et al., 2005), raising the possibility that an ancestral intermediate compartment exists in ERES-Golgi units of *Drosophila* and other organisms.

Another set of unresolved questions in our understanding of the ER-Golgi interface relate to the nature of membrane carriers generated therein. In vitro studies indicate that the COPII machinery directs budding of anterograde vesicles through assembly of a vesicle-coating cage 60-80 nm in diameter (Barlowe et al., 1994; Matsuoka et al., 1998). Reports of COPII vesicles in this size range in cells exist (Bykov et al., 2017; Hughes et al., 2009; Zeuschner et al., 2006), but are rare compared to the numerous studies documenting COPI and Clathrin vesicles (Langhans et al., 2012). This has for long time raised speculations on whether structures different from regular-sized COPII vesicles mediate ER-Golgi transport (Mironov and Beznoussenko, 2019; Robinson et al., 2015). Indeed, many protein cargos are secreted in animal cells that exceed by far the dimensions of a 60-80 nm vesicle (Fromme and Schekman, 2005). Examples of these include collagens, the main components of animal extracellular matrices, for which trimers assemble inside the ER into 300-400 nm long rods (Canty and Kadler, 2005). Conflicting recent studies describe existence (Gorur et al., 2017; Jin et al., 2012; Matsui et al., 2020; Melville et al., 2019; Raote et al., 2017; Santos et al., 2015; Yuan et al., 2018) or absence (McCaughey et al., 2019; Omari et al., 2020) of large megavesicle carriers involved in ER-Golgi collagen transport. To visualize collagen traffic from ERES, these studies prevented collagen trimerization through ascorbate depletion, followed by ascorbate readministration to trigger resumption of transport. This has been shown to cause ER-phagy (Omari et al., 2018), complicating the interpretation of structures formed under these conditions. The alternative to megavesicle carriers as mediators of large cargo transport is direct ER-Golgi connection, for which some evidence exists in budding yeast (Kurokawa et al., 2014) and plants (daSilva et al., 2004). ERES-ERGIC contact has been proposed as a transport mechanism in mammalian cells as well (Malhotra and Erlmann, 2015; Raote and Malhotra, 2021). To date, nonetheless, the existence of each of these types of structures at ERES, namely regular-sized vesicles, megacarriers and/or ER-Golgi connections remains a subject of intense debate.

Yeast has provided for many years a genetically tractable system to research secretion (Barlowe and Miller, 2013; Schekman, 2010). *Drosophila*, however, is increasingly becoming an excellent model to dissect secretion in higher eukaryotes and animals. Most studied proteins involved in secretion have fly homologues, including COPI and COPII components, Rab-GTPases, SNAREs, TRAPP complex, p24 proteins and golgins (Kondylis and Rabouille, 2009). Gene redundancy in these core machineries, however, is very limited when compared to mice or humans. Another advantage of *Drosophila* is the availability of precise genetic tools that allow transgenic protein tagging, forward genetic screening, and loss- and gain-of function experiments. Genetic screenings using these tools have identified conserved new secretory genes (Bard et al., 2006; Ke et al., 2018; Kondylis et al., 2011; Tiwari et al., 2015; Wendler et al., 2010). Among these, Tango1, an ERES-localized transmembrane protein of the metazoan MIA/cTAGE (Melanoma Inhibitory Activity/Cutaneous T-cell lymphoma-associated antiGEn) family, has been shown to function in ERES definition and ERES-Golgi coordination, in addition to postulated roles in collagen transport (Feng et al., 2021; Malhotra and Erlmann, 2015). *Drosophila* is, in addition, a convenient model to investigate the biology of collagen and the extracellular matrix (Pastor-Pareja, 2020). Flies do not possess fibrillar collagens, but produce Collagen IV, the main component of basement membranes (Davis et al., 2019). *Drosophila* Collagen IV, a 450 nm-long heterotrimer, is abundantly present in all fly tissues (Lunstrum et al., 1988). In the larva, the main source of Collagen IV is fat body adipocytes, while other tissues such as imaginal discs, precursors of the adult epidermis, do not produce any (Pastor-Pareja and Xu, 2011). Interestingly, ERES of the Collagen IV-producing fat body are significantly larger than imaginal disc ERES (Liu et al., 2017). However, a detailed comparative study of the ER-Golgi interface in the fat body has not been carried out.

To better understand secretory pathway organization and ER-Golgi traffic, we used structured illumination microscopy (SIM) and focused ion beam scanning electron microscopy (FIB-SEM) to characterize *Drosophil*a ERES-Golgi units. We found in front of ERES a pre-cis-Golgi compartment through which both anterograde and retrograde transport pass. This pre-cis-Golgi is continuous with the rest of the Golgi and not a separate intermediate compartment or a collection of large membrane carriers, for which we found no evidence. In every ERES analyzed through FIB-SEM, however, we found vesicles, as well as tubules extending between ERES and Golgi.

## RESULTS

### Regionalization within *Drosophila* ERES-Golgi units

In order to better understand secretory pathway organization and gain insights into the mechanisms underlying secretory traffic at the ER-Golgi interface, we decided to study the organization of ERES-Golgi units in *Drosophila* through different high-resolution imaging techniques. To do that, we first imaged secretory pathway markers in larval fat body using superresolution 3D-SIM (Three-Dimensional Structured Illumination Microscopy) in ERES-Golgi units of the fat body, the main source of collagen and other extracellular matrix proteins in the fly larva. Consistent, with our previous observations (Liu et al., 2017), Tango1 colocalized with ERES marker Sec16 in multiple irregularly-shaped structures per cell, usually about 1 μm micron in diameter (Figure 1A and B). Golgi Microtubule Associated Protein (GMAP), Mannosidase II (ManII) and Galactosyltransferase (GalT), conserved markers for cis-, mid- and trans-regions of the Golgi apparatus, respectively, showed close but distinct localization within ERES-Golgi units (Figure 1C), showing that, despite its small size, the Golgi element in ERES-Golgi units is regionalized. Furthermore, localization of GMAP and additional cis-Golgi protein GM130 was clearly resolvable, with GM130 present in a position more proximal to ERES than GMAP (Figure 1D). Localization of Grasp65, a third conserved cis-Golgi-associated protein, largely overlapped that of GM130 (Figure 1E). From these data, we conclude that ERES-Golgi units in *Drosophila* contain, in addition to trans-, mid- and cis- regions, a fourth cis-most region that we call pre-cis-Golgi (Figure 1F).

**Figure 1.**
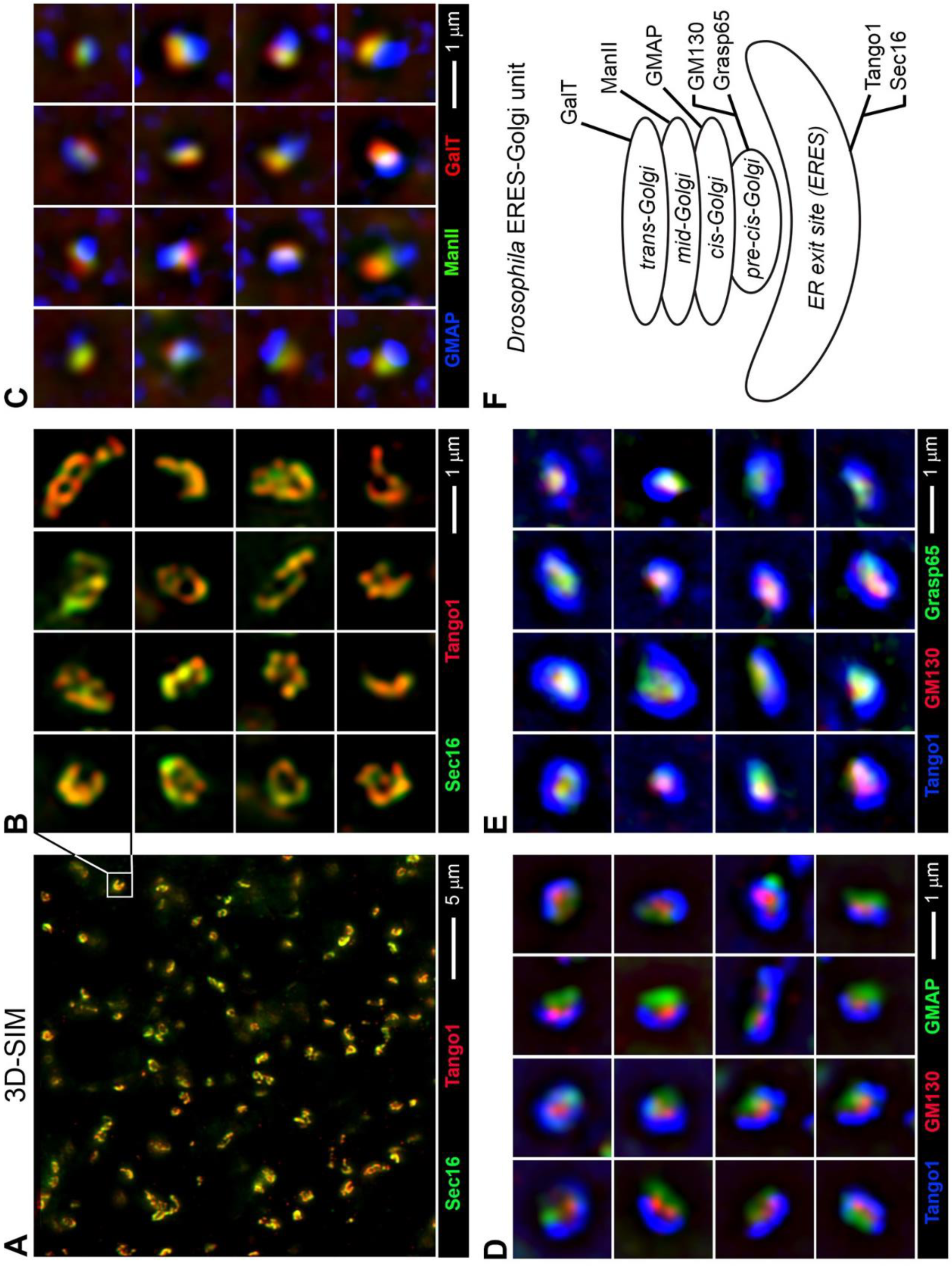
*Drosophila* ERES-Golgi units contain trans-, mid-, cis- and pre-cis-Golgi elements. (A) Super-resolution SIM (Structured Illumination Microscopy) image of third instar larval fat body showing ERES markers Sec16 (*Sec16.sGFP*, green) and Tango1 (antibody staining, red). (B) Magnified view of individual ERES in (A). (C) SIM images of Golgi markers GMAP (antibody staining, blue), ManII (*Cg>ManII.EGFP*, green) and GalT (*Cg>GalT.TagRFP*, red) in fat body. (D) SIM images of ERES marker Tango1 (antibody staining, blue) and Golgi markers GMAP (*Gmap.GFP*, green) and GM130 (antibody staining, red) in fat body. (E) SIM images of ERES marker Tango1 (antibody staining, blue) and Golgi markers Grasp65 (*Cg>Grasp65.GFP*) and GM130 (antibody staining, red) in fat body. (F) Schematic illustration of an ERES-Golgi unit depicting relative localization of ERES and Golgi markers. Besides cis-, mid- and trans-Golgi, a fourth pre-cis-Golgi element can be distinguished.

### COPI and COPII distribution within ERES-Golgi units

The COPII (anterograde) and COPI (retrograde) coat machineries for vesicle budding are essential for secretory transport in eukaryotes (Barlowe and Miller, 2013). To investigate the functional organization of the ER-Golgi interface and the role of the pre-cis-Golgi compartment, we studied the localization within ERES-Golgi units of COPI and COPII proteins. To do that, we created transgenic flies expressing tagged versions of COPI coat protein γCOP, COPI-GTPase Arf1, COPII coat protein Sec13 and COPII-GTPase Sar1. The transgenic versions we constructed of these proteins included dual GFP/APEX2 tags for use in both light and electron microscopy. The tagged proteins, when expressed in the larval fat body, exhibited cytoplasmic localization with clear concentrations in ERES-Golgi units and no apparent effects on cellular health or animal viability. Therefore, we proceeded to study in more detail their localization within ERES-Golgi units through 3D-SIM in combination with additional markers. γCOP signal highlighted structures resembling the ERES element in ERES-Golgi units, and indeed its localization closely paralleled the localization of ERES marker Tango1 (Figure 2A). COPI GTPase Arf1, in contrast, localized to the Golgi apparatus, as shown by colocalization with mid-Golgi marker ManII (Figure 2B). Regarding the localization of COPII proteins, both coat component Sec13 and GTPase Sar1 were found to localize to cis-Golgi; however, Sec13 was found to resemble most closely pre-cis-Golgi Grasp65 (Figure 2C), whereas COPII GTPase Sar1 showed colocalization with proper cis-Golgi marker GMAP (Figure 2D). These results indicate that COPI and COPII components localize to ERES-Golgi unit, where they tend to concentrate in different specific regions (Figure 2E).

**Figure 2.**
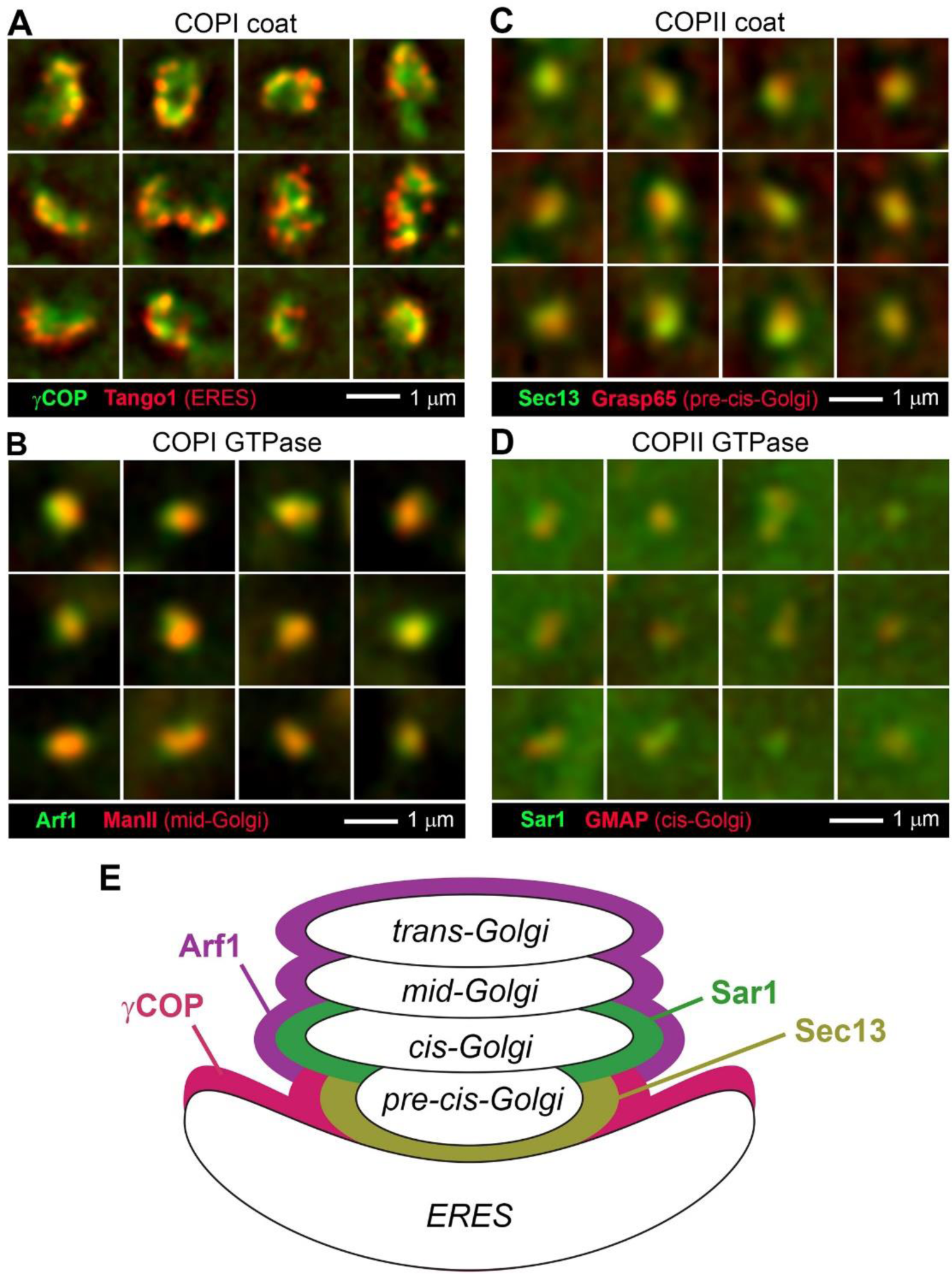
COPI and COPII distribution in ERES-Golgi units imaged through SIM. (A) SIM images showing localization of γCOP (*Cg>γCOP.APEX.GFP*, green) and ERES marker Tango1 (antibody staining, red) in fat body. (B) SIM images showing localization of Arf1 (*Cg>Arf79F.GFP.APEX*, green) and mid-Golgi marker ManII (*Cg>ManII.TagRFP*, red) and ERES marker Tango1 (antibody staining, red) in fat body. (C) SIM images showing localization of Sec13 (*Cg>Sec13.GFP.APEX*, green) and pre-cis-Golgi marker Grasp65 (Cg>Grasp65.RFP, red) in fat body. (D) SIM images showing localization of Sar1 (*Cg>Sar1.GFP.APEX*, green) and cis-Golgi marker GMAP (antibody staining, red) in fat body. (E) Schematic illustration of an ERES-Golgi unit depicting concentration in different regions of COPI and COPII proteins.

To confirm our assessment of COPI and COPII localization and better place them within the ERES-Golgi unit, we imaged these proteins also through their APEX tags, capable of producing upon reaction with DAB dark deposits that can be visualized through transmission electron microscopy (TEM) (Martell et al., 2017). Besides fat body, we analyzed cells of the wing imaginal discs, the larval precursors of the adult wing epidermis. ERES-Golgi units could be recognized in both tissues as discrete ER regions partially surrounding compact clusters of complex membrane elements, as confirmed by the localization of APEX fusions for ERES Tango1 (Figure 3A) and Golgi Grasp65 (Figure 3B), and consistent with our 3D-SIM data and TEM observations of others (Kondylis et al., 2001; Rabouille et al., 1999). In both fat body and imaginal discs, γCOP delineated cup-shaped ERES on their concave sides (Figure 3C). COPI-GTPase Arf1, in contrast, localized to the membrane complex opposed to the ERES (Figure 3D). Consistent with 3D-SIM again, COPII protein Sec13 occupied a central position in the ERES concavity in contact with it (Figure 3E), like γCOP. COPII-GTPase Sar1, finally, concentrated in the Golgi in a position more distal than Sec13 (Figure 3F). In summary, our 3D-SIM and APEX-TEM data show that COPI and COPII coat proteins and GTPases localize within ERES-Golgi units in different but nearby locations.

**Figure 3.**
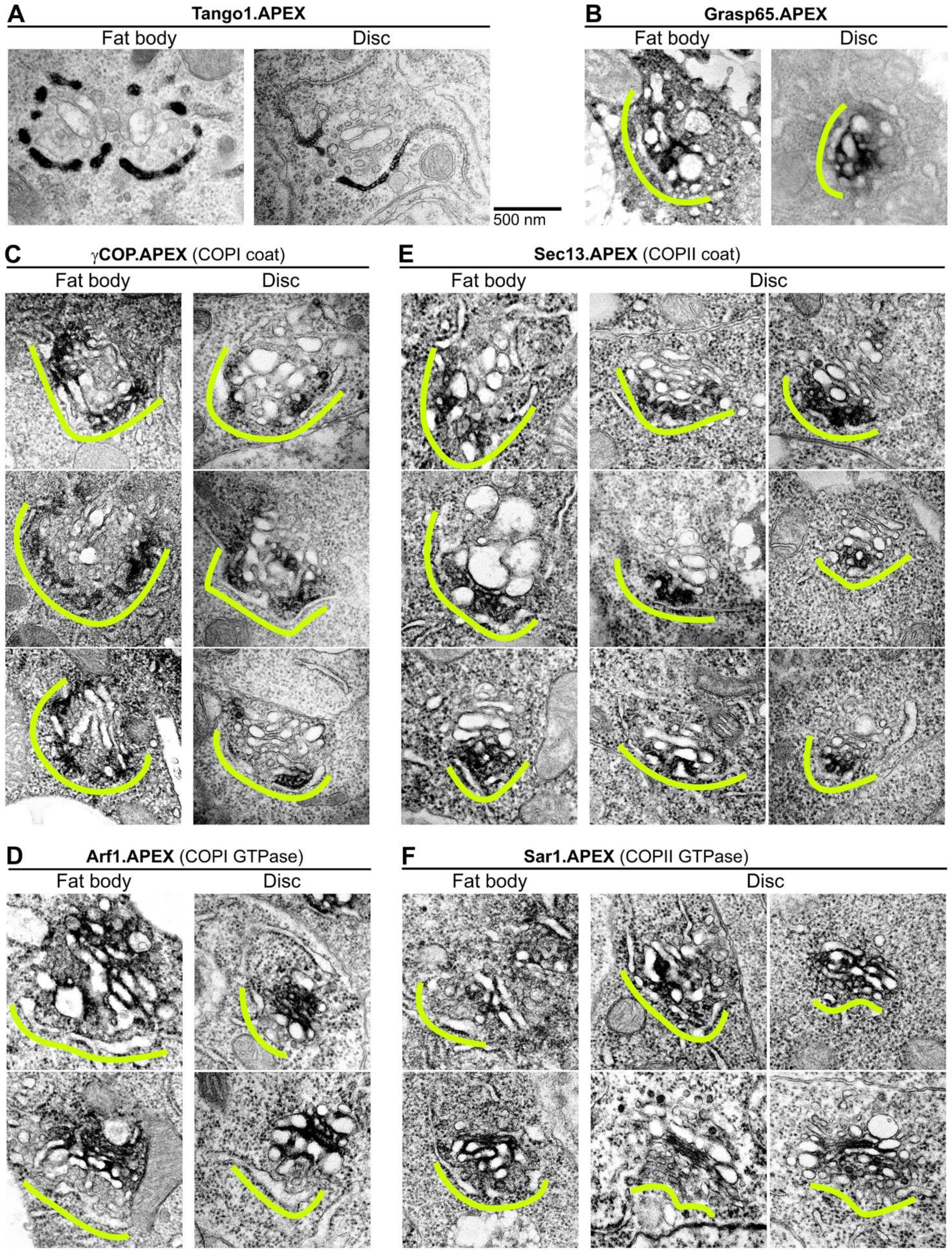
COPI and COPII distribution imaged through APEX-TEM. Transmission electron micrographs from fat body or imaginal disc tissue, as indicated, showing localization within ERES-Golgi units of (A) ERES marker Tango1 (*SP.GFP.APEX.Tango1*), (B) pre-cis-Golgi marker Grasp65 (*Grasp65.GFP.APEX*), (C) COPI coat protein γCOP (*γCOP.APEX.GFP*), (D) COPI GTPase Arf1 (*Arf79F.GFP.APEX*), (E) COPII coat protein Sec13 and (*Sec13.GFP.APEX*), and (F) COPII GTPase Sar1 (*Sar1.GFP.APEX*). Expression of transgenic APEX-tagged proteins was driven by *Act-GAL4* in imaginal discs and by *Cg-GAL4* in fat body. Dark deposits after DAB reaction reveal concentration of APEX-tagged proteins. Yellow lines outline the ERES concavity. All APEX signals are cytoplasmic except for Tango1, tagged in its ER luminal domain.

### Pre-cis-Golgi is involved in both anterograde and retrograde ER-Golgi transport

COPI and COPII coat proteins appear to concentrate in peripheral and central positions of the ERES cup, respectively. To confirm this complementary distribution, reminiscent of recent findings in the yeast *Pichia pastoris* (Roy Chowdhury et al., 2020), we examined ERES-Golgi units in fat body expressing Sec13.GFP and γCOP.RFP simultaneously. This confirmed that COPII is found at the center of ERES, whereas COPI occupies preferentially the periphery of the structure (Figure 4A). Given the differential but contiguous localization of the COPI and COPII coats, suggesting tight coupling of anterograde and retrograde traffic, we next tried to ascertain whether pre-cis-Golgi was involved in anterograde or retrograde transport. In a genetic screening we previously conducted, we had found that Grasp65 was required for efficient general secretion (Ke et al., 2018). As we reported before, Collagen IV.GFP was retained in fat body cells upon Grasp65 knock down (Figure 4B). This retention occurred in the ER, not in the Golgi, as evidenced by the orientation of ERES-Golgi units in contact with regions of intracellular Collagen IV accumulation (Figure 4C). Having established a requirement of Grasp65 in anterograde ER-Golgi traffic, we next examined its possible role in retrograde Golgi-ER transport. To do that, we imaged GFP fused to the ER retention motif KDEL, which targets proteins for retrograde Golgi-ER transport. KDEL.GFP normally localized in the ERES region of ERES-Golgi units, suggesting efficient recycling of KDEL.GFP from the Golgi back to ERES. Knock down of Grasp65, in contrast, produced localization of KDEL.GFP in front of ERES, indicating that KDEL.GFP was not efficiently trafficked from the Golgi to the ER (Figure 4D). These results show that the pre-cis-Golgi is involved in both anterograde and retrograde transport.

**Figure 4.**
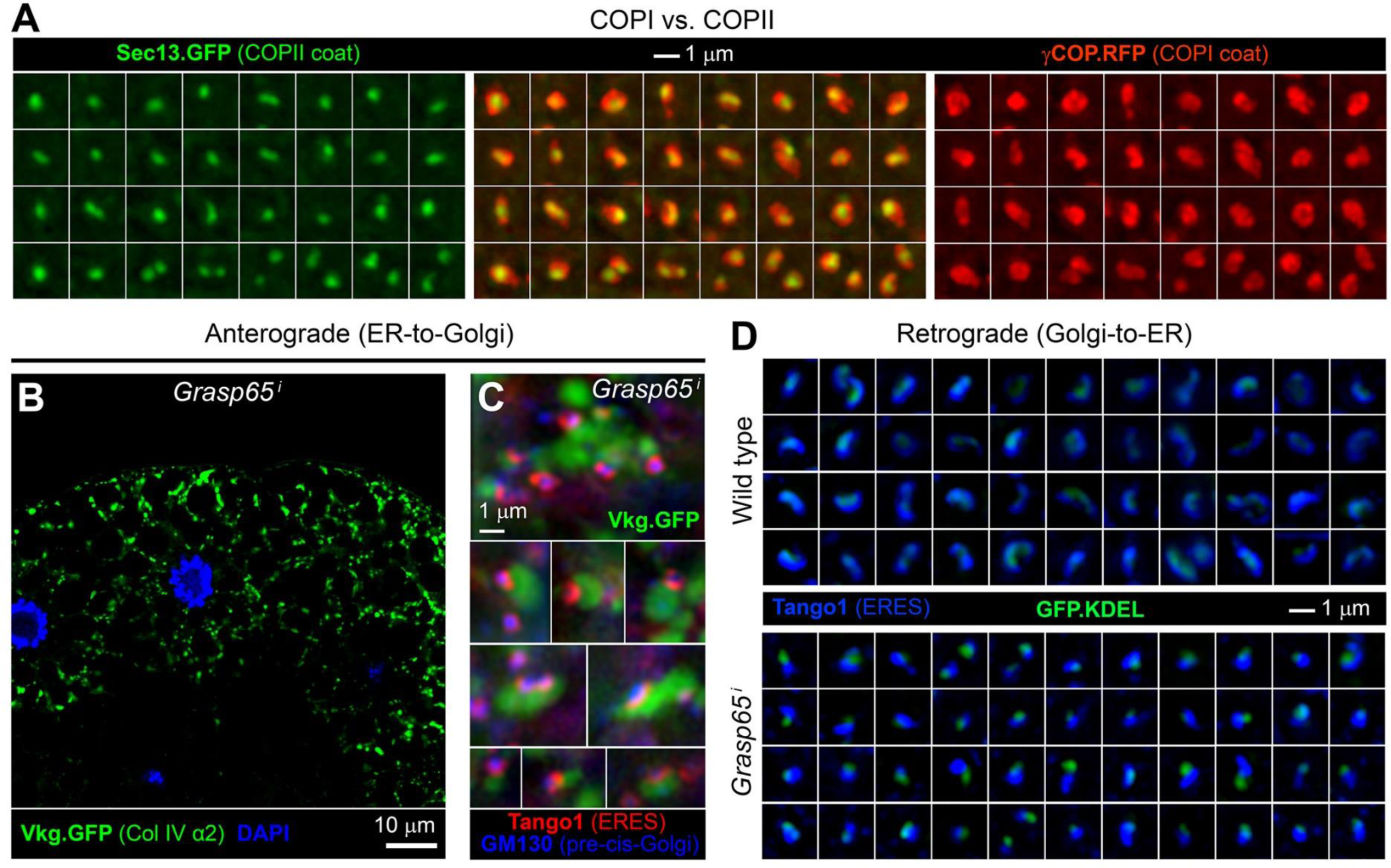
Pre-cis-Golgi is involved in both anterograde and retrograde transport. (A) SIM images of fat body ERES showing localization of COPII coat protein Sec13 (*Cg>Sec13.GFP.APEX*, left panels, green) and COPI coat protein γCOP (*Cg>γCOP.mRFP*, right panels, red) in a complementary center/periphery distribution (center panels, green and red merged). (B) Confocal image of fat body cells showing intracellular retention of Collagen IV (*vkg.GFP*, green) upon Grasp65 knock down (*Cg>Grasp65^i^*). Nuclei stained with DAPI (blue). (C) SIM images of Collagen IV (*vkg.GFP*, green) retained inside fat body cells upon Grasp65 knock down (*Cg>Grasp65^i^*). The tissue has been stained with anti-Tango1 (ERES, red) and anti-GM130 (Golgi, blue). The orientation of ERES-Golgi units with respect to retained collagen indicates ER retention. (D) SIM images of retrograde cargo GFP.KDEL (*Cg>GFP.KDEL*, green) distribution in relation to ERES (anti-Tango1, blue) in wild type (top) and *Cg>Grasp65^i^* (bottom) fat body. Accumulation of GFP.KDEL in front of ERES upon Grasp65 knock down indicates Golgi retention.

### FIB-SEM analysis of ERES-Golgi units in fat body and imaginal disc cells

Intrigued by the nature of the ERES/pre-cis-Golgi interface, which our TEM sections could not clarify, we decided to investigate ERES-Golgi units using FIB-SEM. This technique allows serial sectioning and electron microscopy imaging of plastic-embedded tissues for 3D reconstruction of subcellular structures with high resolution in the z axis (Narayan and Subramaniam, 2015). Through FIB-SEM, we imaged volumes of fat body and of wing imaginal disc tissue with a z resolution of 20 nm (Figure 5A and B, and Suppl. Video 1). ERES-Golgi units were readily recognizable on the basis of morphology alone. Also recognizable were contact sites between ER and other organelles (Figure S1). With data acquired from two samples of each tissue, and using Dragonfly software, we constructed 3D models of 15 fat body and 15 wing disc ERES-Golgi units. (Figure 5C and D, and Suppl. Video 2). In addition, we reconstructed 15 imaginal disc ERES-Golgi units from one sample of imaginal wing discs expressing Tango1 (see next Results section and Figure S2). From the analysis and comparison of the morphological traits of fat body and wing disc ERES-Golgi units, general characteristics of these structures became apparent. ERES appeared in concave regions of ribosome-covered ER formed by convergence of fenestrated ER sheets (Figure 5E and F). In front of these, the Golgi apparatus, despite tremendous complexity and high tubulation, appeared in all units analyzed as a single, fully continuous structure (Figure 5C-F). On the trans side of the Golgi, saccular and tubular elements could be observed, including some identifiable as lysosome-related degradative bodies. Within each ERES-Golgi unit, ERES proper were usually discontinuous, archipelago-like collections of ER membrane patches devoid of ribosomes (Figure 5G and H). Between ERES and Golgi, numerous isolated vesicles could be observed, as well as ERES-Golgi tubes (Figure 5G and H). We proceeded next to analyze ERES, Golgi and intervening membrane structures in more detail.

**Figure 5.**
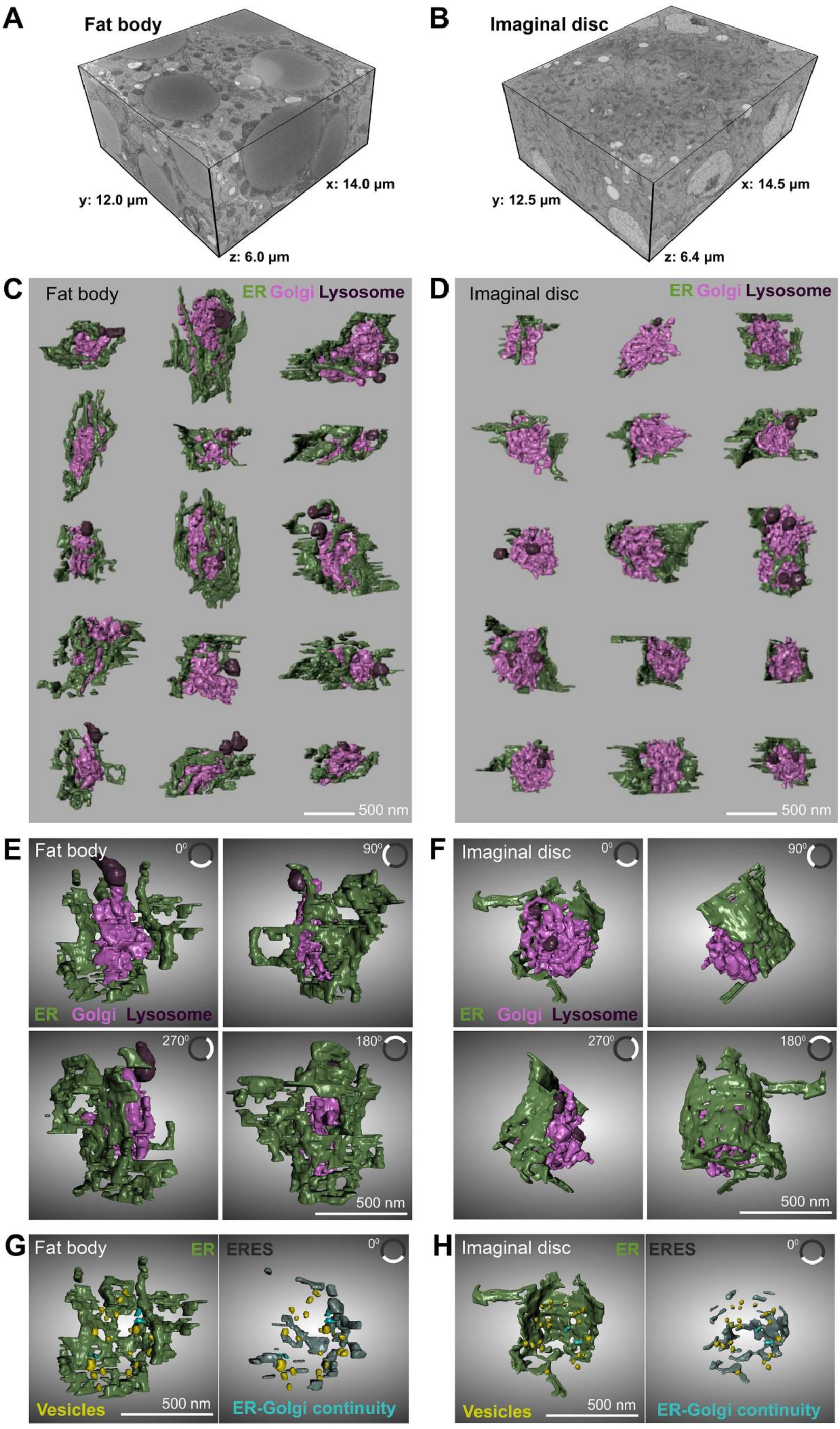
FIB-SEM analysis reveals a single continuous Golgi compartment per ERES-Golgi unit. (A, B) Dimensions of two FIB-SEM volumes obtained from wild type third-instar larval fat body (A) and wing imaginal disc (B) tissues. See also Suppl. Video 1. (C, D) 3D models of 15 fat body (C) and 15 imaginal disc (D) ERES-Golgi units reconstructed from FIB-SEM data. Different colors indicate ER (green), Golgi (light purple) and lysosome-related degradative structures (dark purple). See also Suppl. Video 2. (E, F) Horizontally rotated views of single fat body (E) and imaginal disc (F) ERES-Golgi units, colored as in C and D. The rotation angle of each view is provided in the upper right corner of each panel. See also Suppl. Video 3 and 4. (G, H) Frontal views of the ERES concavity in ERES-Golgi units from fat body (G, same unit as E) and imaginal disc (H, same unit as F). ER (left, green), ERES (right, grey), vesicles (yellow) and ER-Golgi continuities (blue) are represented.

### Tango1 expression increases ERES size

We performed measurements of ERES and Golgi size on ERES-Golgi units reconstructed from our FIB-SEM data. ERES, defined as ER regions devoid of ribosomes on their Golgi-facing side (Figure 6A), were larger in fat body than in wing imaginal disc cells (Figure 6B, C and F), consistent with our previous observations using SIM (Liu et al., 2017). Overexpression of Tango1 in the imaginal disc increased the size of this ER region devoid of ribosomes on one side (Figure 6D and F), indicating an increase in the size of ERES. In contrast with the difference in ERES size, Golgi size was not significantly different between fat body and imaginal disc tissues (Figure 6G). Correlation between ERES size and Golgi size within each ERES-Golgi unit, consistently, was weak (Figure 6H). Golgi morphology, however, was markedly different between fat body and imaginal discs, as cisternae stacked in a cis-trans direction were distinguishable in imaginal disc ERES-Golgi units (4.9±0.6 cisternal levels, n=15) (Figure 6E). In contrast, cisternal stacking was not obvious in fat body ERES-Golgi units, which displayed a more globular, amorphous morphology. Consistent with this, the ratio surface/volume was higher in imaginal disc ERES-Golgi units (Figure 6I). In all, these results, comparing fat body and imaginal discs, suggest that ERES size can vary across cell types depending on the level of Tango1 expression, while Golgi size, despite differences in the degree of cisternal organization, varies less.

**Figure 6.**
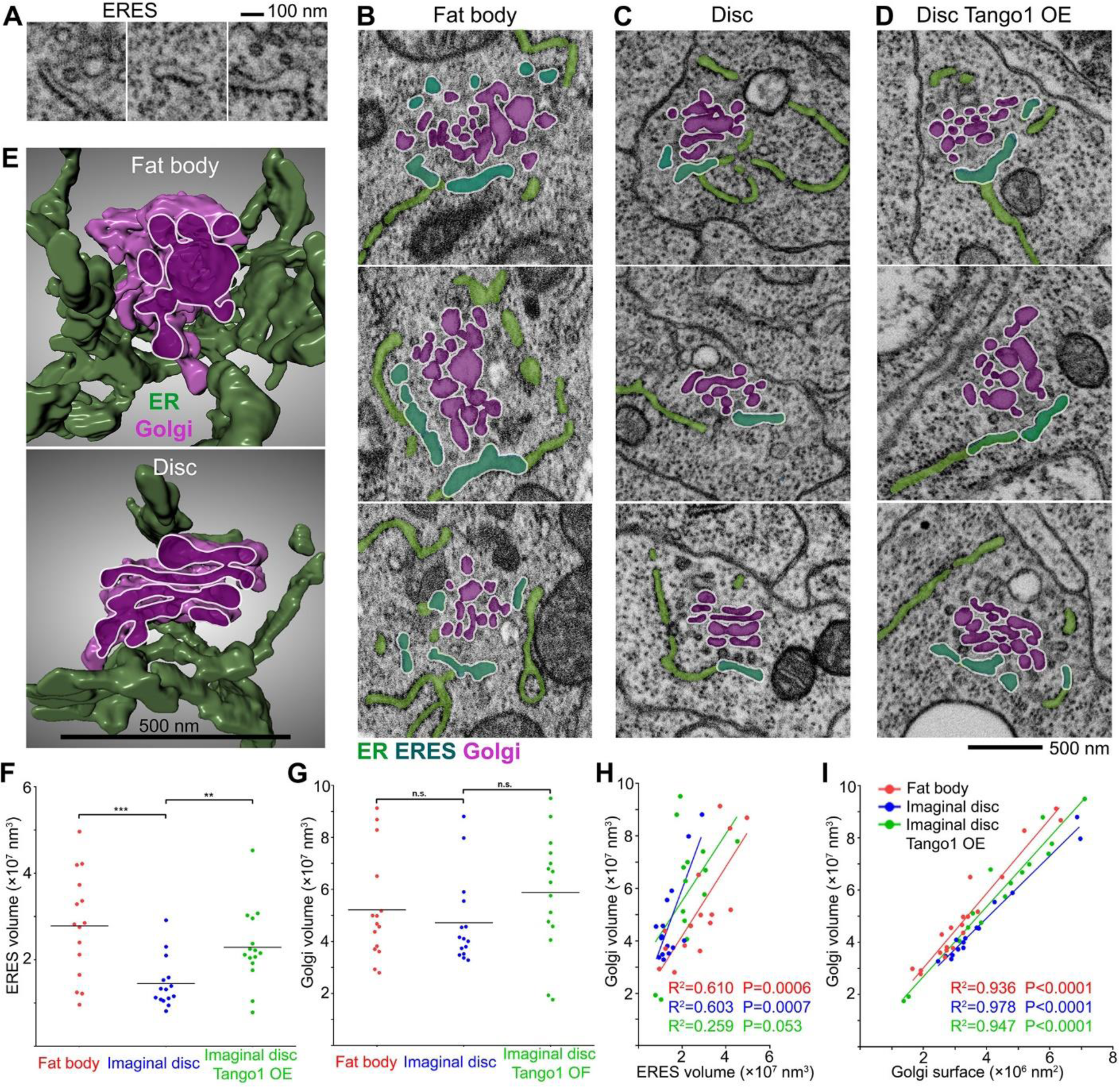
Tango1 expression increases ERES size. (A) FIB-SEM images featuring examples of ER devoid of ribosomes on its Golgi-facing side (ERES). (B-D) FIB-SEM images of ERES-Golgi units from wild type fat body (B), wild type wing imaginal disc (C) and wing imaginal disc overexpressing Tango1 (D, *Act>SP.GFP.Tango1*). Superimposed colors indicate ER (light green), ERES (dark green) and Golgi (purple). (E) 3D reconstructions of ERES-Golgi units from wild type fat body (top panel) and imaginal disc (bottom panel). ER (green) and Golgi (light purple) are depicted. A cis-trans section through the Golgi is shown (dark purple). (F, G) Quantification of ERES (F) and Golgi (G) volume as measured in 3D models of ERES-Golgi units. Each dot represents a single ERES-Golgi unit from wild type fat body (red), wild type imaginal disc (blue) and Tango1-overexpressing imaginal disc (Act>SP.GFP.Tango1, green). Horizontal lines represent mean values. Significance was determined using two-tailed t tests. Fat body ERES (n=15) vs. imaginal disc ERES (n=15), p=0.001 (***). Imaginal disc ERES (n=15) vs. imaginal disc Tango1 OE ERES (n=15), p=0.0048 (**). Fat body Golgi (n=15) vs. imaginal disc Golgi (n=15), p=0.4728 (not significant). Imaginal disc Golgi (n=15) vs. imaginal disc Tango1 OE Golgi (n=15), p=0.1879 (not significant). (H, I) Correlation Golgi volume vs. Golgi surface (H) and Golgi volume vs ERES volume (I), as measured in 3D models of ERES-Golgi units. Each dot represents a single ERES-Golgi unit (n=15 in each group) from wild type fat body (red), wild type imaginal disc (blue) and Tango1-overexpressing imaginal disc (*Act>SP.GFP.Tango1*, green). P value and R^2^ were determined using linear regression tests.

### ERES-Golgi vesicles and pearled tubes

Between the Golgi apparatus and the ER, we found numerous vesicles in ERES-Golgi units of both fat body (Figure 7A) and imaginal disc tissue (Figure 7F). In total, we identified 286 vesicles in ERES-Golgi units of fat body (Figure 7B) and 363 in those of imaginal discs (Figure 7G). The number of vesicles in each ERES-Golgi unit ranged from 8 to 33 in fat body (Figure 7C) and from 14 to 47 in imaginal disc tissue (Figure 7H). Besides vesicles, also visible were omega-shaped buds, emerging from both ERES and Golgi (Figure 7D, E, I and J). We next analyzed vesicle size and localization within the ERES-Golgi unit of these vesicles. To do this, we measured the diameter of these vesicles, and found that in both fat body and imaginal disc tissue the distribution of vesicle sizes showed two peaks at 52 nm and 64 nm (Figure 7B and G), suggesting that these corresponded to two different populations of vesicles. Furthermore, when the position within ERES-Golgi units of vesicles was mapped with a cutoff distinguishing vesicles larger and smaller than 58 nm in diameter, their distribution was reminiscent of the relative COPI center/COPII periphery distribution observed in our SIM data (Figure 7K and L). In 15 ERES-Golgi units of imaginal disc cells overexpressing Tango1, finally, we identified 369 vesicles, which showed a similar two-peaked diameter distribution (Figure S2). In all, our data show that vesicles are abundant in ERES-Golgi units in *Drosophila*.

**Figure 7.**
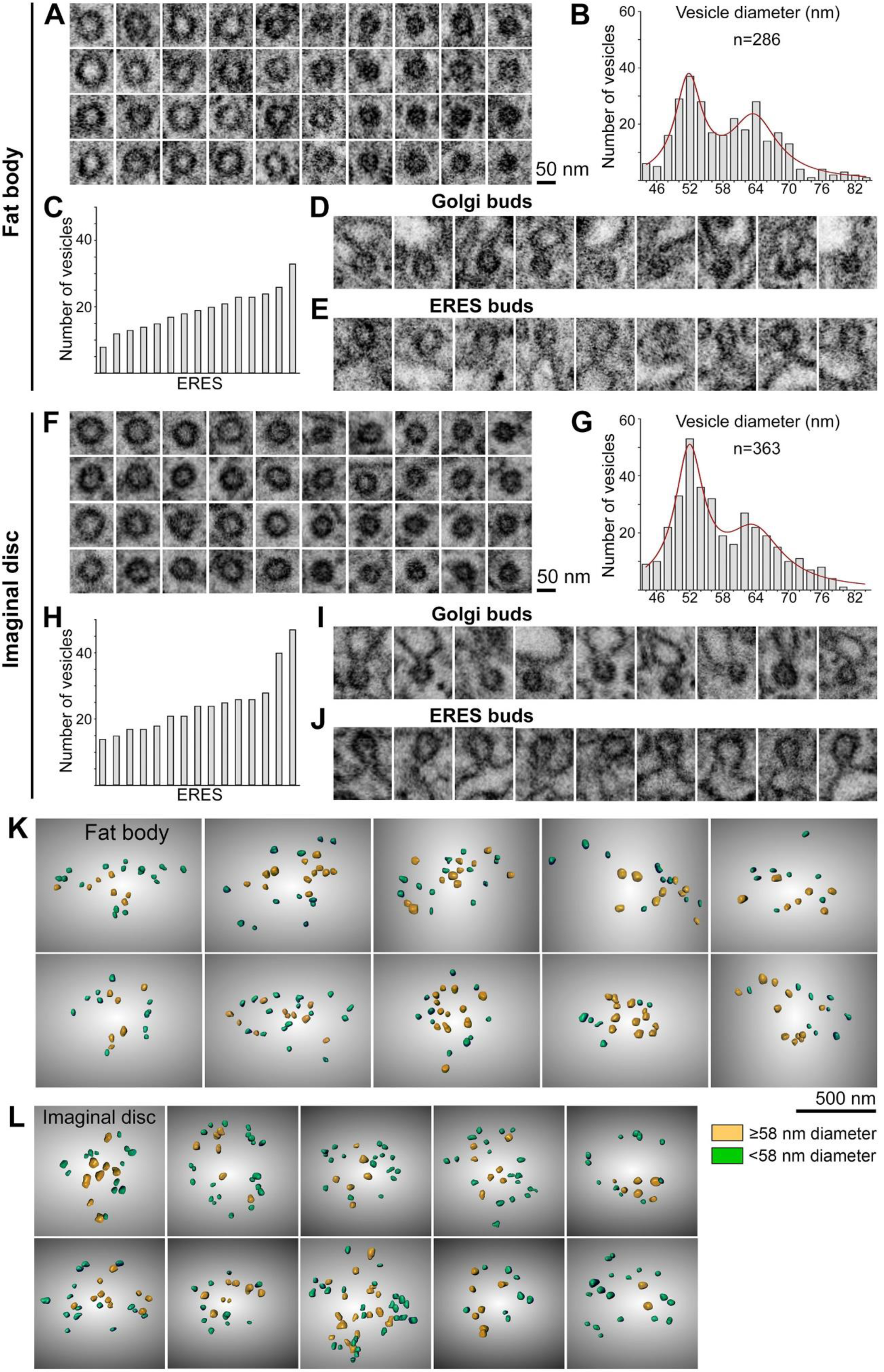
Analysis of vesicles in the ER-Golgi interface. (A, F) FIB-SEM images of vesicles found in ERES-Golgi units of wild type fat body (A) and wing imaginal disc (F) tissues. (B, G) Frequency distribution of vesicle diameters in the ER-Golgi interface of 15 fat body (B) and 15 imaginal disc (G) ERES-Golgi units. The red line fits the distribution to a sum of Lorentzians curve. Diameter of a vesicle was measured on FIB-SEM images directly, not on 3D models, as the maximum diameter in any FIB-SEM section of that vesicle (Z-resolution of our data was 20 nm). (C, H) Number of vesicles in 15 fat body (C) and 15 imaginal disc (H) ERES-Golgi units. Each column represents an individual ERES. (D, E, I, J) FIB-SEM images of buds found in Golgi (D, I) and ERES (E, J) of fat body (D, E) and imaginal disc (I, J) tissues. (K, L) Frontal views of the ERES concavity from 3D models of fat body (K) and imaginal disc (L) ERES-Golgi units showing the spatial distribution of vesicles. Vesicles larger and smaller than 58 nm in diameter are represented in yellow and green, respectively.

In addition to vesicles and buds, our FIB-SEM data showed tubular connections between the ERES and Golgi. Connections spanned distances of about 100 nm between ERES proper and Golgi and were never tubes with parallel walls, but had in all instances a pearled shape (Figure 8A and B). In a majority of cases (>80%), their shape was one-beaded, resembling a vesicle connected to the ERES and Golgi compartments by narrow necks (Figure 8C). Besides one-beaded tubes, we found some examples of two- and three-beaded tubes (Figure 8B and C). Instances of two-beaded and three-beaded tubes extending from ERES without contacting Golgi could be identified as well (Figure 8D and E). In total, we found at least one connection in each ERES-Golgi unit we reconstructed, with a maximum of seven in one imaginal disc unit (Figure 8F). Same as buds, tubules associated in all cases with ERES proper (Figure 8G and H). In summary, our imaging of ERES-Golgi units found abundant regular-sized vesicles and pearled tubes at the ER-Golgi interface. In contrast, in none of the 45 ERES-Golgi units analyzed by FIB-SEM we found evidence of larger megacarrier vesicles, free tubular elements or saccules capable of carrying a cargo like collagen from ERES to Golgi.

**Figure 8.**
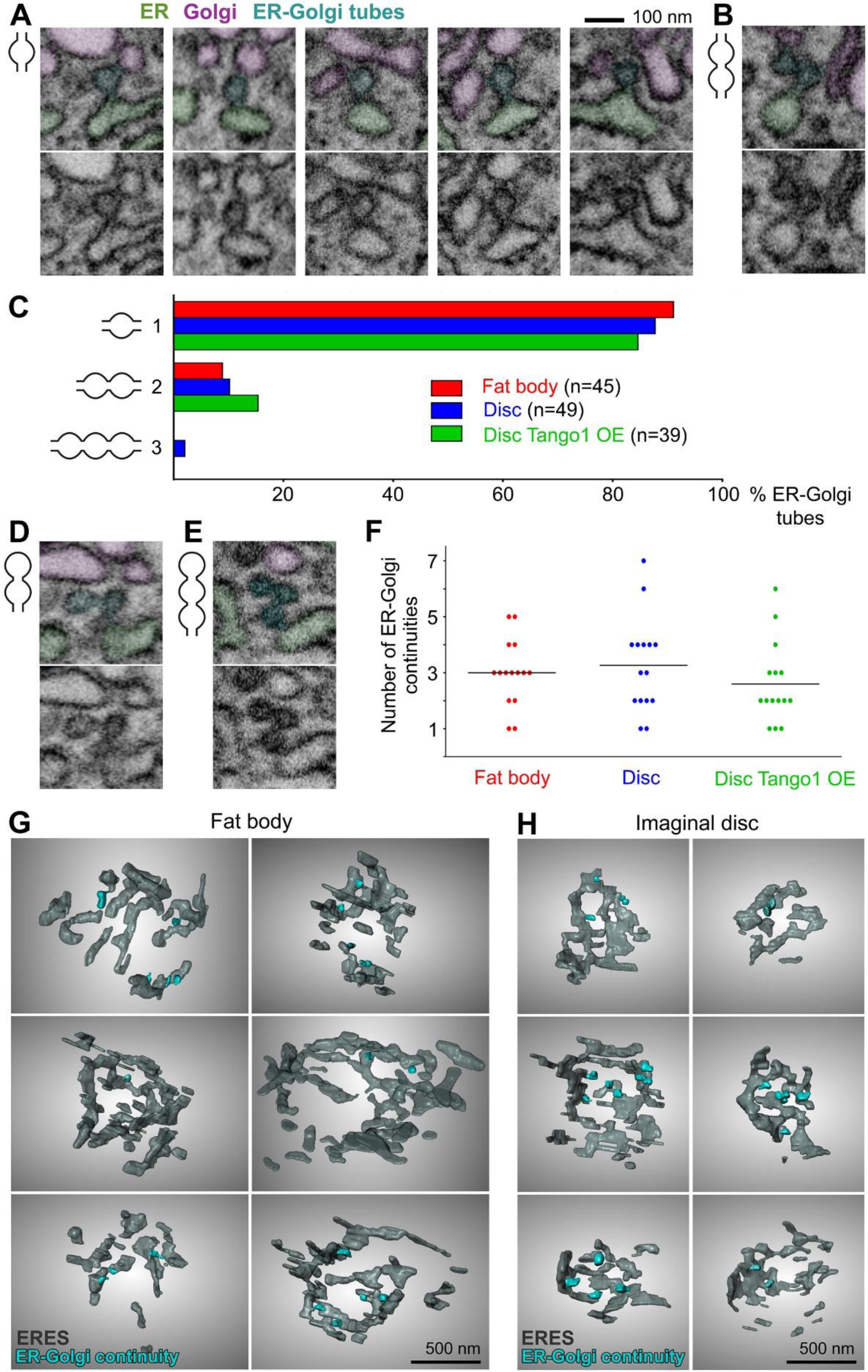
ERES-Golgi connection through pearled tubular continuities. (A, B) FIB-SEM images of one-beaded and two-beaded (B) tubes connecting ERES and Golgi. In top panels, ER, Golgi and connecting tubes are pseudocolored in green, purple and cyan, respectively. Bottom panels show the original image. (C) Percentage of one-beaded, two-beaded and three-beaded ERES-Golgi continuities from the total observed in 15 ERES-Golgi units from fat body (n=45 continuities), 15 from imaginal disc (n=49) and 15 from Tango1-overexpressing imaginal discs (n=39). (D, E) FIB-SEM images of two-beaded (D) and three-beaded (E) tubes extending from ERES. In top panels, ER, Golgi and ERES tubes are pseudocolored in green, purple and cyan, respectively. Bottom panels show the original image. (F) Number of ERES-Golgi continuities in each ERES-Golgi unit analyzed. Each dot represents an ERES-Golgi unit (n=15 in each group). Horizontal lines mark mean values. (G, H) Frontal views of the ERES concavity from 3D models of fat body (G) and imaginal disc (H) ERES-Golgi units showing the position of ERES-Golgi continuities (cyan) in relation to ERES (grey).

## DISCUSSION

### Architecture of *Drosophila* ERES-Golgi units

We used high resolution imaging techniques, transgenic protein tagging, and chosen loss- and gain-of-function conditions to characterize *Drosophila* ERES-Golgi units. Importantly, we used FIB-SEM data to reconstruct the 3D architecture of 45 ERES-Golgi units from fat body and imaginal disc tissues. Although very heterogeneous in shape, intra- and inter- tissue comparisons revealed general features. In both fat body and imaginal discs, ERES are located in concave regions of ER. These are not flat cups, but consist of convergent ER sheets which are similar to the rest of the fly ER, although more fenestrated and tubulated. On this ER concavity, a collection of areas devoid of ribosomes conforms the ERES proper. Tango1 overexpression in imaginal discs increased the size of this ribosome-devoid area, consistent with a role of Tango1 in defining ERES (Liu et al., 2017; Reynolds et al., 2019; Rios-Barrera et al., 2017). Golgi lies at a distance of about 100 nm, giving rise together with ERES to a compact structure, recognizable in both TEM and FIB-SEM samples. It has been proposed that ERES are phase-separated, membrane-less organelles that behave like liquid droplets (Gallo et al., 2020; Hanna et al., 2018). Viscosity due to concentration of traffic regulators between ERES and Golgi must be high, indeed. However, our characterization shows *Drosophila* ERES-Golgi units as compact membrane assemblages. While liquid-liquid phase separation may be crucial for aspects of ERES establishment and function (Maeda et al., 2020; Zhang and Rabouille, 2019), their maintenance may be better explained by limited diffusion imposed by the enclosing membranes and classical tethering of those membranes by proteins like Tango1, Grasp65, golgins and Rab1. Indeed, in plants, there is evidence of physically solid attachment between ERES and Golgi (Sparkes et al., 2009).

Despite its amorphous appearance and weak cisternal organization, the markers we studied indicate that the Golgi is regionalized into cis-, mid- and trans-elements. Additionally, markers characteristic of cis-Golgi could be resolved in two locations defined by GMAP (cis-Golgi) and Grasp65/GM130 (pre-cis-Golgi). FIB-SEM reconstructions, nonetheless, invariably showed the Golgi as a single continuous structure. Therefore, the pre-cis-Golgi region is not a separate cisterna or tubular cluster, but an integral part of the Golgi apparatus. We conclude, thus, that an intermediate compartment between ERES and Golgi does not exist in *Drosophila* as a separate entity. Furthermore, our characterization of Grasp65 loss shows that the pre-cis-Golgi region where it localizes is involved in both anterograde and retrograde transport. This, together with the fact that Grasp65 and GM130 are found in vertebrate ERGIC (Marra et al., 2001), suggests that the ERGIC is not just functionally equivalent, but also evolutionarily homologous to the fly pre-cis-Golgi. This comparative view would support a status for the ERGIC as a structural constituent of the Golgi rather than a transient collection of carriers, placing the actual ER-Golgi interface directly in front of vertebrate ERES, as in other eukaryotes. Consistent with this, a recent study in human cells found that during ER-Golgi transport COPII components remain in the vicinity of ERES, cargo traveling from there in COPII-uncoated, Rab1-dependent carriers (Westrate et al., 2020).

Unlike ERES, which were larger in fat body cells and increased in size with Tango1 overexpression, Golgi size was not significantly different across samples and only weakly correlated with ERES size, suggesting that partially autonomous organizing mechanisms act in ERES and Golgi despite their intimate relation. A difference was patent however in Golgi morphology, as cisternal organization was more distinct in disc cells than in fat body. In mammalian cells, increased secretory load is known to enlarge ERES on one hand (Farhan et al., 2008), and induce higher cisternal connectivity on the other (Trucco et al., 2004). Fat body adipocytes are highly secretory cells producing not just Collagen IV and other extracellular matrix proteins, but also large amounts of serum proteins, clotting factors and antibacterial peptides. Differential secretory activity, therefore, may underlie morphological divergence between fat body and imaginal disc ERES-Golgi units. *Drosophila*, hence, could be an excellent model for future studies of self-organizational properties, physiological adaptation and scaling relations across the ER-Golgi interface with the help of recently developed inducible cargos (Casler et al., 2020).

While in the budding yeast *Saccharomyces cerevisiae* anterograde ER-Golgi and retrograde Golgi-ER transport seem to take place in different ER regions (Schröter et al., 2016), in the fission yeast *Pichia pastoris* they are tightly coupled at ERES, with COPII and COPI displaying a center/periphery relative distribution when visualized at high resolution (Roy Chowdhury et al., 2020). Our *Drosophila* data show now the same COPII-center/COPI-periphery relation in an animal. Broad colocalization of SEC16 with Tip20 homologue RINT1 in U2OS cells suggested closeness between ERES and ERAS as well in human cells (Roy Chowdhury et al., 2020). It would be interesting to know next whether in vertebrates this relative center/periphery relation is conserved at the same fine scale and how this relates to the existence of the ERGIC.

### Membrane structures mediating ER-Golgi transport

In our FIB-SEM data, abundant vesicles can be observed between ERES and Golgi in all 45 ERES-Golgi units we modeled. In many instances, presence of a coat in vesicles and buds is discernible from the darker signal outlining them. The resolution of our imaging, however, does not allow determination of the type of coat for each vesicle. Nonetheless, the distribution of vesicle sizes, with two peaks around 52 and 64 nm in diameter, suggests the presence of two different populations, which might correspond to COPI and COPII vesicles, respectively. Consistent with this would be also the spatial arrangement of the vesicles, correlating with the COPII-center/COPI-periphery distribution observed with SIM and APEX-TEM. In addition to vesicles, we observed tubular connections between Golgi and ERES. These were not straight tubes, but pearled tubulations, the majority of them consisting of a single vesicle-sized bead, suggesting that vesicle budding machineries must be involved in their formation. The simplest explanation for this kind structure would be a budding vesicle meeting the opposite compartment before excision. However, the few two-beaded and three-beaded tubes we observed are difficult to explain through this mechanism. Multibudded tubules have been observed protruding from ERES in cultured human cells (Bannykh et al., 1996) and their formation can be induced by COPII on liposomes (Bacia et al., 2011). Alternatively, buds from Golgi and ERES could meet to form a tube. Because our study is not dynamic, we have no insight into the duration of connections. However, it is worth noting that their prolonged maintenance should result in bidirectional flow, difficult to reconcile with transport directionality. If, on the contrary, the tubes are short-lived intermediates preceding excision from the donor compartment, directional transport may still be achieved (Mironov and Beznoussenko, 2019).

Tubular elements have been associated with all steps of secretion (Martínez-Menárguez, 2013; Mironov et al., 2003; Polishchuk et al., 2009; Robinson et al., 2015; Simpson et al., 2006). Our data suggests that vesicles, because of their large numbers, are the predominant form of ER-Golgi exchange. Nonetheless, our results may have important implications for the transport of collagen and other cargos that cannot fit into these vesicles. Collagen-specific factors creating enlarged carriers have been postulated. Our data in *Drosophila*, documenting 1018 vesicles and 133 ER-Golgi tubes, found no evidence for megavesicles. This is despite the fact that we analyzed fat body, a cell type producing Collagen IV, the major collagen in flies. We therefore conclude that, absent megavesicles, tubular continuities are the only option left capable of transporting such proteins. We also observed tubular continuities in ERES of imaginal discs, which do not produce collagen, indicating that these are not specific of collagen-secreting cells. We can speculate that in vertebrates, where Golgi and ERES are distant, an evolutionary parsimonious translation of our findings would support that tubular connections between ERES and ERGIC, rather than megavesicles, mediate ER export of large proteins (Malhotra and Erlmann, 2015; McCaughey et al., 2019). If true, reports of larger vesicles transporting collagen to the Golgi, if not artefactual (Omari et al., 2020), could correspond to COPII-uncoated ERGIC membranes (Westrate et al., 2020).

## MATERIALS AND METHODS

### *Drosophila* strains

Standard fly husbandry techniques and genetic methodologies, including balancers and dominant markers, were used to assess segregation of transgenes in the progeny of crosses, construct intermediate lines and obtain flies of the required genotypes for each experiment (Roote and Prokop, 2013). Flies were cultured at 25°C in all experiments. The GAL4-UAS binary expression system (Brand and Perrimon, 1993) was used to drive expression of UAS transgenes under temporal and spatial control of transgenic GAL4 drivers *Cg-GAL4* (fat body), *BM-40-SPARC-GAL4* (fat body) and *Act5C-FO-GAL4* (imaginal discs). Stable insertion of transgenic UAS constructs was achieved through standard P-element transposon transgenesis (Rubin and Spradling, 1982). Detailed genotypes in each experiment are provided in Table S1. The following strains were used:

*w^1118^* (used as wild type; Bloomington Drosophila Stock Center, 3605)

*w; Cg-GAL4* (Bloomington Drosophila Stock Center, 7011)

*y w; Act5C-FO-GAL4 / TM6B,Tb* (Bloomington Drosophila Stock Center, 3954)

*y w; Sec16.sGFP^fTRG.1259^* (Vienna Drosophila Resource Center, 318329)

*w Gmap^KM0132^.GFP* (Kyoto Drosophila Genomics and Genetics Resources, 109702)

*w; UAS-GalT.TagRFP; TM2 / TM6B,Tb* (Bloomington Drosophila Stock Center, 65251)

*w; UAS-ManII.EGFP; TM2 / TM6B,Tb* (Bloomington Drosophila Stock Center, 65248)

*w; UAS-ManII.TagRFP* (Bloomington Drosophila Stock Center, 65249)

*w; UAS-Grasp65.GFP* (Bloomington Drosophila Stock Center, 8507)

*w; UAS-Grasp65.RFP* (this study)

*y w; Kr^If-1^ / CyO*; *UAS-γCOP.mRFP* (Bloomington Drosophila Stock Center, 29714)

*w; UAS-γCOP.APEX.GFP* (this study)

*w; UAS-Arf79F.GFP.APEX* (this study) *w; UAS-Sec13.GFP.APEX* (this study) *w; UAS-Sar1.GFP.APEX* (this study)

*w; UAS-SP.GFP.Tango1* (Liu et al., 2017).

*w; UAS-SP.GFP.APEX.Tango1* (this study)

*w; UAS-Grasp65.GFP.APEX* (this study)

*y sc v sev; UAS-Grasp65 ^HMC05584^.RNAi* (Bloomington Drosophila Stock Center, 64565)

*w; vkg^G454^.GFP / CyO*; *BM-40-SPARC-GAL4 UAS-Dcr2 / TM6B* (Zang et al., 2015)

*w; UAS.GFP.KDEL* (Bloomington Drosophila Stock Center, 30906)

### UAS-Sec13.GFP.APEX, UAS-Sar1.GFP.APEX, UAS-Arf1.GFP.APEX and UAS-Grasp65.GFP.APEX

Gateway destination vector pTGW (UASt-GFP-Gateway cassette, Drosophila Carnegie Vector collection) was modified into pTWGA (UASt-Gateway cassette-GFP-APEX). For that, the APEX sequence was PCR-amplified from plasmid pcDNA3 APEX2-NES (Addgene, cat # 49386) with primers attNheIAPEX-F and attSpeIAPEX-R adding att sites. The resulting fragment was purified through gel extraction (Magen HiPure Gel Pure DNA Mini kit, cat # D2111-03,) and cloned into vector pDONR221 (Thermo Fisher Scientific, cat # 12536017) with Gateway^TM^ BP Clonase^TM^ II Enzyme Mix (Thermo Fisher Scientific, cat # 11789020) to produce a pDONR221-APEX entry clone. From there, the APEX sequence was recombined into pTGW through Gateway LR recombination using LR Clonase^TM^ II Plus enzyme (Thermo Fisher Scientific, cat # 12538120) to obtain pTGA. Because this vector now lacked a Gateway recombination cassette, the cassette was added back to pTGA using an Xbal restriction site at the 5’ of GFP. To do this, first, the Gateway casette sequence was PCR-amplified from pTGW with primers sooXbaIGate-F and sooXbaIGate-R adding XbaI restriction sites and soo sites. Gel purified PCR product soo-XbaI-Gateway-XbaI-soo was then cloned into pTGA linearized by XbaI digestion (New England Biolabs, cat#R0145S) using SoSoo mix reagent (Trelief^TM^ SoSoo Cloning Kit, TSINGKE, cat # TSV-S1) to generate vector pTWGA.

To produce each individual APEX.GFP construct, the coding sequence of each gene was amplified from whole larva cDNA using PrimeScript RT-PCR Kit (Takara, cat # RR014-A). The resulting products were recombined into pDONR221 and transferred to the pTWGA vector to finally produced the desired plasmids. Primers were: attSec13-F, attSec13-R; attGrasp65-F, attGrasp65-R; attSar1-F, attSar1-R; attArf79F-F, attArf79F-R.

### UAS-SP.GFP.APEX.Tango1

The Gateway cassette sequence was added to pT-SP-G (Liu et al., 2017) to obtain pTSGW (UASt-Signal peptide of Tango1-GFP-Gateway cassette). For that, the cassette sequence was amplified from pTGW with primers SpeIgate-F and XhoIgate-R’, adding SpeI and XhoI restriction sites and the resulting fragment was inserted into pT-SP-G through SpeI and XhoI double enzyme digestion and T4 ligation, producing pTSGW destination vector, into which APEX.Tango1 sequence from pDONR221-APEX.Tango1 was later transferred through Gateway LR recombination.

To obtain pDONR221-APEX.Tango1, the GFP sequence in pT-SP-G-Tango1 (Liu et al., 2017) was removed through NheI (New England Biolabs, cat # R3131L) and SpeI (New England Biolabs, cat # R3133L) restriction. In its place, we introduced the APEX sequence, PCR-amplified from pcDNA3 APEX2-NES with primers NheIAPEX-F’ and SpeIAPEX-R, which added NheI and SpeI restriction sites for subsequent double enzyme digestion and ligation with T4 DNA ligase (New England Biolabs, cat # M0202L) to obtain pTSA-Tango1. From pTSA-Tango1, the APEX.Tango1 sequence was amplified with primers attAPEXTango1-F and attAPEXTango1-R, adding att sites. The resulting att-flanked APEX.Tango1 fragment was cloned into pDONR221 through Gateway BP recombination to obtain pDONR221-APEX.Tango1 entry clone.

### UAS-γCOP.APEX.GFP

The coding sequence of γCOP was obtained by RT-PCR from whole larva cDNA with primers γCOP-F and γCOP-APEX-R. The APEX sequence was PCR-amplified from pcDNA3 APEX2-NES with primers γCOP-APEX-F and APEX-R. Overlap PCR was used to link γCOP and APEX sequences into γCOP.APEX. γCOP.APEX was then PCR-amplified with primers attγCOP-F and attAPEX-R adding att sites and cloned into pDONR221 through Gateway BP recombination to obtain pDONR221-γCOP.APEX. From there, the γCOP.APEX sequence was transferred into pTWG (UASt-Gateway cassette-GFP, Drosophila Carnegie Vector collection) through Gateway LR recombination.

### UAS-Grasp65.RFP

Grasp65 was amplified with primers adding att sites as above, recombined into plasmid pDONR221 using Gateway BP recombination to obtain pDONR221-Grasp65, and from there transferred to pTWR (UASt-Gateway cassette-RFP, Drosophila Carnegie Vector collection) using Gateway LR recombination.

### SIM and confocal imaging

Tissue samples were predissected in PBS by turning them inside out with fine tip forceps, fixed in PBS containing 4% PFA (paraformaldehyde, Sinopharm Chemical Reagent, cat # 80096692), washed in PBS (3 × 10 min), dissected from the carcass and mounted on a glass slide with a drop of DAPI-Vectashield (Vector Laboratories, cat # H-1200). SIM image stacks (z-steps of 0.24 μm) were acquired with a Nikon A1 N-SIM STORM microscope equipped with a CFI Apo SR TIRF 100× oil (NA 1.49) objective and an Andor Technology EMCCD camera (iXON DU-897 X-9255). Laser lines at 488, 561 and 640 nm were used for excitation. SIM image reconstructions were performed with NIS-Elements software (Nikon). Images are maximum intensity projections of two to five confocal sections. Confocal images of *Grasp65^i^* fat body (Figure 4C) were acquired in a ZEISS LSM780 microscope equipped with a 100× oil Plan-Apochromat objective (NA 1.4).

### Immunohistochemistry

The following primary antibodies were used: Guinea pig anti-Tango1 (1:1,000) (Lerner et al., 2013), rabbit anti-GM130 (1:500, Abcam, cat # ab30637) and goat anti-Gmap (1:500) (Riedel et al., 2016). Secondary antibodies were Goat Anti-Guinea Pig IgG (Rhodamine conjugated, Jackson ImmunoResearch, cat # 106025003; Alexa Fluor 647 conjugated, Jackson ImmunoResearch, cat # 106605003), Alexa Fluor 555 Donkey Anti-Rabbit IgG (Invitrogen, cat # 1945911) and Alexa Fluor 555 Donkey Anti-Goat IgG (Abcam, cat # ab150130) respectively. Antibody stainings were perfomed using standard procedures for larval tissues. Briefly, larvae were predissected in PBS, fixed in PBS containing 4% PFA (paraformaldehyde, Sinopharm Chemical Reagent, cat # 80096692), washed in PBS (3 × 10 min), blocked in PBT-BSA [PBS containing 0.1% Triton X-100 detergent (Sigma-Aldrich, cat # T8787), 1% BSA (Zhongkekeao, cat # 201903A28), and 250 mM NaCl (Amresco, cat # 0805C384)], incubated overnight with primary antibody in PBT-BSA in 4°C, washed in PBT-BSA (3 × 20 min), incubated for 2 h with secondary antibody in PBT-BSA at RT, washed in PBT-BSA (3 × 20 min) and PBS (3 × 10 min). Tissues were finally dissected and mounted on a glass slide with a DAPI-Vectashield (Vector Laboratories, cat # H-1200).

**APEX-TEM**

Third instar larvae were predissected by turning them inside out with fine tip forceps in fixation solution containing 2.5% glutaraldehyde (TED PELLA, cat # 18426), 2% paraformaldehyde (Alfa Aesar, cat # 43368) and 0.1 M PB (Na2HPO4 (Alfa Aesar, cat # A11817), NaH_2_PO_4_ (Amresco, cat # 0571-500G), pH 7.2). Prefixation was conducted at RT for 2h in the same solution. After this, predissected larvae were washed in 0.1 M PB (3 X 7 min, RT). During the last wash, 30% H_2_O_2_ (Guoyao, cat # 10011218) was quickly mixed in DAB solution to a 0.03% v/v concentration. To prepare the DAB solution, DAB (3,3’-Diaminobenzidine, Sigma, cat # D56637-1G) was freshly dissolved in 0.1 M PB (pH 7.2) to a 0.5 mg/ml concentration and kept at 4°C avoiding light. The larval carcasses were incubated in the H_2_O_2_/DAB mixture on a glass depression well at 4°C for 5-10 min, gently shaking in the dark. After this, the carcasses were transferred to 0.1 M PB and post-fixed in a mixture of 1% osmic acid (Tedpellco, cat # 018456) and potassium ferrocyanide (1.5% w/v, Sigma, cat # 60299-100G-F) at RT avoiding light. Postfixation times were 30 min for fat body 30 min and 20 min for imaginal disc tissues. Samples were then washed with MiliQ H_2_O (5 X 7 min, RT), incubated with 1% uranyl acetate (Merck, cat # 8473) in dark at RT for 1 h or overnight at 4°C, washed with MiliQ H_2_O (5 X 7 min, RT), and dehydrated in a series of 5 min washes on ice with prechilled 30%, 50%, 70%, 90% and 100% (twice) ethanol (Tongguang Jingxi Huagong, cat # 104021), acetone/ethanol (1:1) and 100% acetone (Tongguang Jingxi Huagong, cat # 105003). Infiltration was conducted at RT with a mixture of acetone and resin 1:1 for 1.5 h, 1:2 for 3 h and 1:3 overnight. The next day, carcasses were immersed in resin (3 X 3 h and then overnight, RT) consisting of v/v 48% SPI-PON 812 (Epoxy Resin Monomer, SPI-CHEM, cat # 02659-AB), 16% DDSA (Dodecenyl Succinic Anhydride, SPI-CHEM, cat # 02827-AF), 34% NMA (Nadic Methyl Anhydride, SPI-CHEM, cat # 02828-AF) and 2% BDMA (N,N-Dimethylbenzylamine, SPI-CHEM, cat # 02821-CA). The desired tissues were then dissected from the carcasses and placed in block molds filled with resin for hardening at 60°C during 48 h. 70 nm ultrathin section from the hardened blocks were cut on a Leica EM UC7 ultramicrotome using an Ultra 45° diamond knife (DiATOME, Knives No # MS17502) and imaged in a Hitachi H-7650B electron microscope.

### FIB-SEM

Resin blocks containing embedded sample tissues were obtained as above. Blocks were trimmed to expose tissues, adhered to a 45/90° screw type holder (φ12.7 mm*22.8 mm, Zhongxingbairui, ZB-Y1811), and coated with Gold using a HITACHI E-1010 ion sputter coater for 120 s. FIB-SEM imaging was performed in a FEI Helios NanoLab G3 dual beam microscope system equipped with ETD, TLC, and ICD cameras (ThermoFisher Scientific). A 0.6 μm protective Pt coat was applied to the region of interest through gas injection prior to FIB milling and SEM imaging. For milling slices, an ion beam current of 0.43 nA at 30 kV acceleration voltage was used for milling slices at a step size of 20 nm. The parameters for SEM imaging were: 0.4 nA beam current, 2 kV acceleration voltage, 2 mm working distance, 8 µs dwell time, 4 nm pixel size and 4096 X 3536 pixel count. TLD and ICD cameras collected backscattered signal for imaging. The imaging software was AutoSlice and View G3 1.7.2 (FEI).

Images acquired by FIB-SEM were imported into Dragonfly (Object Research Systems) using Dragonfly Image Loader and aligned through the SSD method in the Slide Registration panel. For segmentation, different ROIs were created in the ROI tools panel. Each organelle or membrane element was manually segmented as an individual ROI with the ROI Painter round brush tool in 2D mode. After segmentation in sections, ROIs were exported and saved as object files. Objects were then converted into 3D meshes using the export box in the ROI Tools panel to create meshes. Meshes were smoothed 4-6 times and observed in 3D scene mode as solid, fully opaque objects. Observing angles were adjusted manually or set with the flip/rotate tools in the main panel. For volume and surface measurements, values for each object were read in the information panel and recorded. The diameter of each vesicle, visible in 2-4 continuous sections, was measured on its largest xy section in 2D mode by using the Ruler tool in the annotation panel. Movie Maker tools within Dragonfly were used to create movies of rotating ERES-Golgi units.

### Statistical analysis

Statistical analysis and graphical representations were performed using GraphPad Prism. Unpaired two-tailed t tests were performed to test significance of differences in ERES and Golgi volume, and proportion of vesicle classes among ERES-Golgi units of different tissues. Linear regression was performed to test correlation between Golgi surface and volume, as well as ERES and Golgi volume. Vesicle diameter frequency distributions were fit to a sum of Lorentzians curve.

## ACKNOWLEDGEMENTS

We wish to thank Manfred Auer for initial suggestions on Dragonfly use. We also thank Ying Liu and the Tsinghua Center for Protein Research and Technology (Ying Li and Xiaomin Li) for technical help. This work was funded by grants 91854207, 31771600, 31750410689 and 31550110204 (to J.C.P.-P) and 31701248 (to M.L.) from the Natural Science Foundation of China.

## SUPPLEMENTAL MATERIALS

**Fig. S1.**
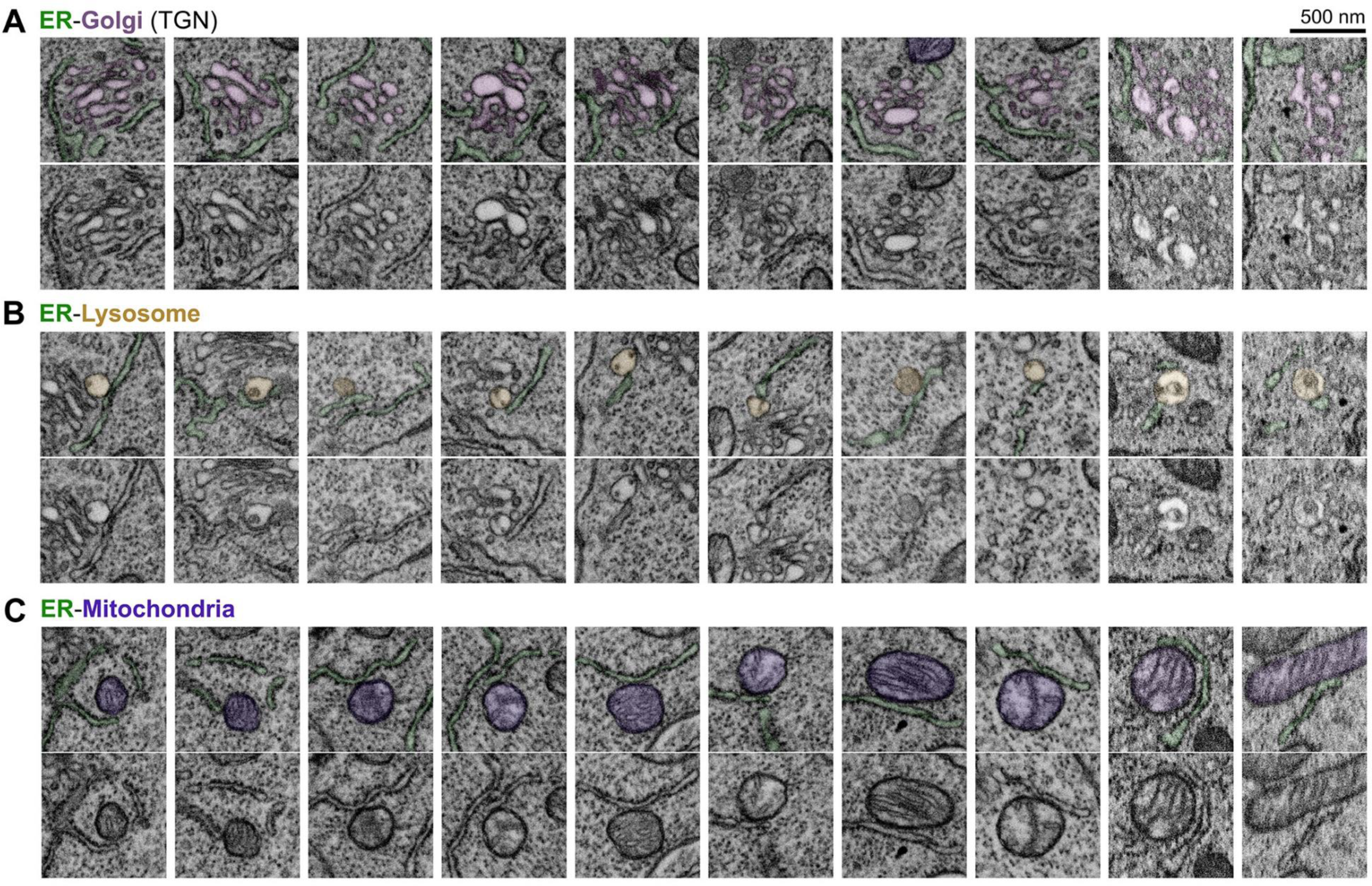
ER contacts with other organelles. FIB-SEM images exemplifying ER-TGN (A), ER-Lysosome (B) and ER-Mitochondria (C) contact sites. In top panels, ER, Golgi, mitochondria and lysosome are pseudocolored in green, light purple, dark purple and yellow, respectively. Bottom panels show the original image.

**Fig. S2.**
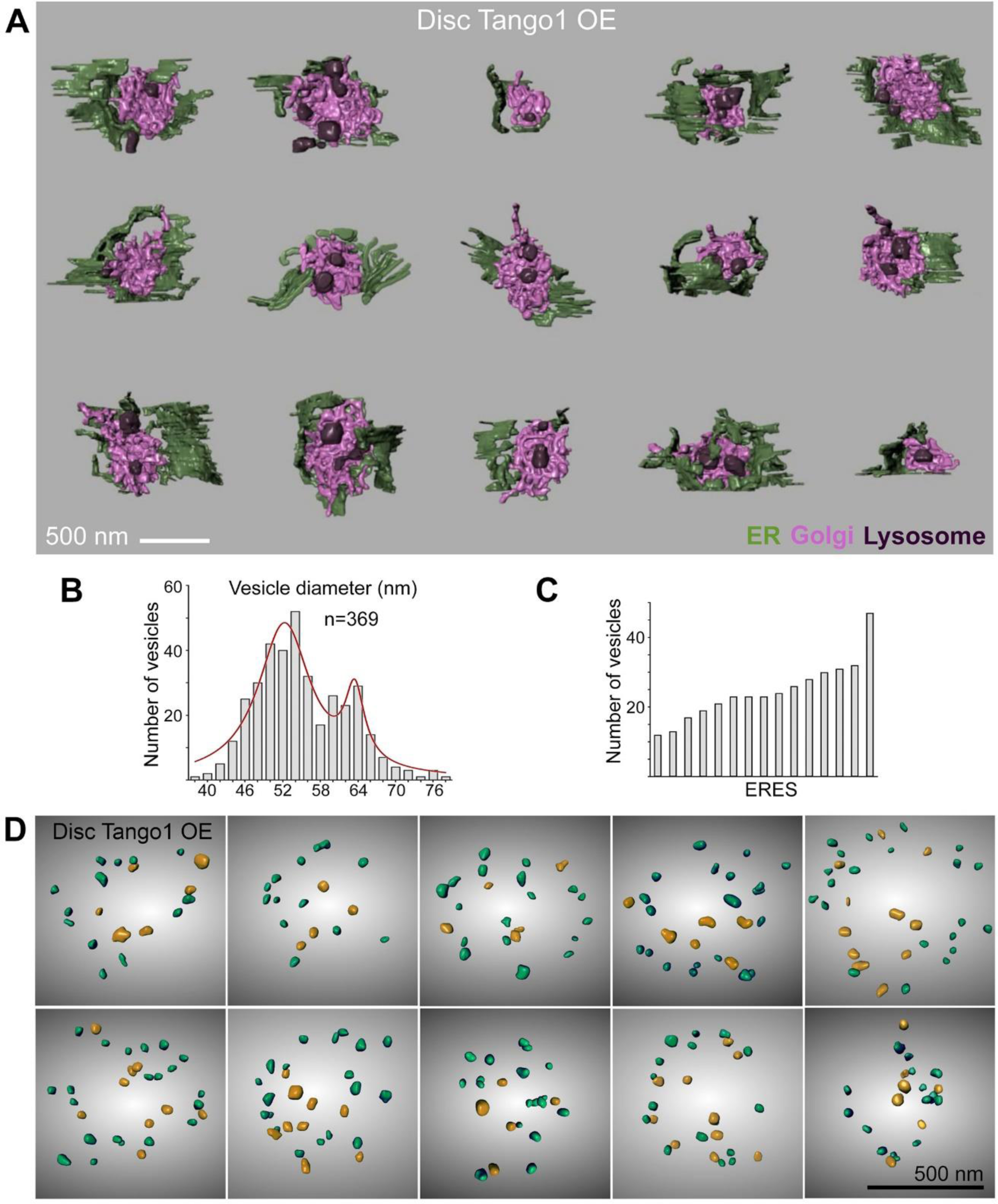
FIB-SEM analysis of ERES-Golgi units in Tango1-overexpressing imaginal disc cells. (A) 3D models of 15 ERES-Golgi units reconstructed from FIB-SEM data of Tango1-overexpressing wing imaginal disc tissue (*Act>SP.GFP.Tango1*). Different colors represent ER (green), Golgi (light purple) and lysosome-related degradative structures (dark purple). (B) Frequency distribution of vesicle diameters in the ER-Golgi interface of ERES-Golgi units from Tango1-overexpressing imaginal disc tissue. The red line fits the distribution to a sum of Lorentzians curve. (C) Number of vesicles in 15 ERES-Golgi units of Tango1-overexpressing disc cells. Each column represents an individual ERES. (D) Frontal views of the ERES concavity from 3D models of ERES-Golgi units of Tango1-overexpressing disc tissue. Vesicles larger and smaller than 58 nm in diameter are represented in yellow and green, respectively.

**Table S1. Detailed genotypes** Genotypes of animals in all experiments, listed by figure.

**Table S2.** Primers

PCR primers used in generation of constructs described in Materials and Methods.

Video S1. FIB-SEM volumes

Serial FIB-SEM registered images of wild type fat body and wild type wing imaginal disc samples, walking through the whole tissue volume in Z-direction.

Video S2. 3D reconstruction of ERES-Golgi units from FIB-SEM data

3D reconstructions of 15 wild type fat body, 15 wild type wing imaginal disc and 15 Tango1 overexpression wing imaginal disc (A*ct>SP.GFP.Tango1*). ER is shown in green, Golgi in light purple and lysosomes in dark purple.

Video S3. 3D model of a fat body ERES-Golgi unit

3D reconstruction of an ERES-Golgi unit from wild type fat body. ER is shown in green, Golgi in light purple, lysosomes in dark purple, vesicles in yellow and ERES-Golgi tubules in blue.

Video S4. 3D model of a wing imaginal disc ERES-Golgi unit

3D reconstruction of an ERES-Golgi unit from wild type wing imaginal disc tissue. ER is shown in green, Golgi in light purple, lysosomes in dark purple, vesicles in yellow and ERES-Golgi tubules in blue.

